# Modeling suggests SARS-CoV-2 rebound after nirmatrelvir-ritonavir treatment is driven by target cell preservation coupled with incomplete viral clearance

**DOI:** 10.1101/2024.09.13.613000

**Authors:** Tin Phan, Ruy M. Ribeiro, Gregory E. Edelstein, Julie Boucau, Rockib Uddin, Caitlin Marino, May Y. Liew, Mamadou Barry, Manish C. Choudhary, Dessie Tien, Karry Su, Zahra Reynolds, Yijia Li, Shruti Sagar, Tammy D. Vyas, Yumeko Kawano, Jeffrey A. Sparks, Sarah P. Hammond, Zachary Wallace, Jatin M. Vyas, Jonathan Z. Li, Mark J. Siedner, Amy K. Barczak, Jacob E. Lemieux, Alan S. Perelson

## Abstract

In a subset of SARS-CoV-2 infected individuals treated with the oral antiviral nirmatrelvir-ritonavir, the virus rebounds following treatment. The mechanisms driving this rebound are not well understood. We used a mathematical model to describe the longitudinal viral load dynamics of 51 individuals treated with nirmatrelvir-ritonavir, 20 of whom rebounded. Target cell preservation, either by a robust innate immune response or initiation of nirmatrelvir-ritonavir near the time of symptom onset, coupled with incomplete viral clearance, appear to be the main factors leading to viral rebound. Moreover, the occurrence of viral rebound is likely influenced by time of treatment initiation relative to the progression of the infection, with earlier treatments leading to a higher chance of rebound. Finally, our model demonstrates that extending the course of nirmatrelvir-ritonavir treatment, in particular to a 10-day regimen, may greatly diminish the risk for rebound in people with mild-to-moderate COVID-19 and who are at high risk of progression to severe disease. Altogether, our results suggest that in some individuals, a standard 5-day course of nirmatrelvir-ritonavir starting around the time of symptom onset may not completely eliminate the virus. Thus, after treatment ends, the virus can rebound if an effective adaptive immune response has not fully developed. These findings on the role of target cell preservation and incomplete viral clearance also offer a possible explanation for viral rebounds following other antiviral treatments for SARS-CoV-2.

**Importance:** Nirmatrelvir-ritonavir is an effective treatment for SARS-CoV-2. In a subset of individuals treated with nirmatrelvir-ritonavir, the initial reduction in viral load is followed by viral rebound once treatment is stopped. We show the timing of treatment initiation with nirmatrelvir-ritonavir may influence the risk of viral rebound. Nirmatrelvir-ritonavir stops viral growth and preserves target cells but may not lead to full clearance of the virus. Thus, once treatment ends, if an effective adaptive immune response has not adequately developed, the remaining virus can lead to rebound. Our results provide insights into the mechanisms of rebound and can help develop better treatment strategies to minimize this possibility.

## Introduction

A 5-day course of nirmatrelvir-ritonavir (N-R) is recommended for individuals who test positive for SARS-CoV-2 with mild-to-moderate symptoms and a high risk of progression to severe disease [1]. Treatment with two doses (300 mg of nirmatrelvir and 100 mg of ritonavir) per day is suggested to be initiated as soon as possible and within 5 days of symptom onset.

Nirmatrelvir is a protease inhibitor, targeting the SARS-CoV-2 main protease 3-chymotrypsin– like cysteine protease enzyme (3CLpro), blocking SARS-CoV-2 replication. Ritonavir reduces the liver catabolism of nirmatrelvir and thus prolongs the half-life of nirmatrelvir [1]. While N-R substantially reduces the risk of progression to severe COVID-19 and can shorten the duration of disease in high-risk individuals [2–4], in some cases, viral rebound and recurring symptoms occur after the 5-day treatment course, including in individuals who have been vaccinated and/or boosted [5,6]. Some individuals with viral rebound are reported to have culturable virus up to 16 days after the initial diagnosis [6,7], thus, potential transmission to close contacts during the rebound period is a concern [5]. Although virus resistant to N-R *in vitro* [8,9] and treatment-emergent 3CLpro substitutions *in vivo* [1,10] have been observed, viral rebound in the case of N-R *in vivo* does not seem to be caused by the emergence of drug resistant mutants [5–7,11–14].

However, two immunocompromised individuals, who were treated with extended duration of N-R in combinations with other treatments, experienced viral rebound associated with resistant mutations E166 A/V and L50F in the NSP5 region where 3CLpro is located [15,16].

The precise proportion of individuals treated with N-R that exhibit viral rebound is unclear, and estimates could vary based on a range of factors, including the definition used to classify rebound and viral characteristics. For example, in the N-R phase 3 clinical trial, EPIC-HR, the fraction of individuals with viral rebound (positive PCR test) and recurring symptom was 1-2% [17]. However, this study was limited by the relatively infrequent viral RNA measurements after the completion of N-R. Other studies have reported rebound in 0.8 – 27% of N-R treated individuals [6,18–22]. Viral rebound has also been described in untreated individuals [23,24], but often at a lower frequency compared to N-R treated individuals regardless of rebound definition [6,17,19,20,22,25,26].

Previously, we analyzed the data presented in Charness et al. [5], where quantitative PCR is available for three individuals who experienced viral and symptom rebound after taking N-R. In all three individuals, no resistance mutations in the gene encoding the protease targeted by nirmatrelvir (3CLpro) developed during treatment and there was no evidence of reinfection by a different variant. The viral dynamic models in our study adequately captured the viral rebound dynamics in all three individuals [27]. One hypothesis we tested was that a 5-day N-R treatment course started near the time of symptom onset reduces the depletion of target cells but does not fully eliminate the virus, thus allowing the virus to rebound once treatment is stopped. The occurrence of viral rebound was shown to be sensitive to model parameters, especially the time therapy is started and the time adaptive immune response begins to emerge. This suggested that a delay in the treatment initiation can lower the chance of rebound. However, our results were only supported by a limited data set comprised of three individuals [27].

Here, we expand upon this previous study using data from an ongoing observational cohort study, including 51 individuals treated with N-R, 20 of whom were classified as having viral rebound per the definition by Edelstein et al. [6] (additional details in Data). Our model accurately captured the viral dynamics of all 51 individuals and provides further evidence that target cell preservation plays a central role in the occurrence of large amplitude viral rebounds. Our model predicts that target cell preservation was achieved by a robust innate immune response or by early treatment. As treatment only stops viral replication but does not directly eliminate existing virus, residual virus may remain after treatment has ended and can infect the remaining target cells and rebound. While we use N-R as a case study, our theory can also explain the viral rebound observed after treatment with molnupiravir [21], another oral antiviral with FDA emergency use authorization, simnotrelvir/ritonavir [28], a protease inhibitor that also targets the SARS-CoV-2 main protease 3CLpro but has a shorter half-life [29] than nirmatrelvir, and VV116 or mindeudesivir [30], an inhibitor of the viral RNA-dependent RNA polymerase that is not inferior to N-R in reducing time to recovery [31].

## Results

### Model of viral dynamics in the upper respiratory tract

We used an extension of a viral dynamic model that has been applied to study SARS-CoV-2 infection dynamics [32–35]. In this model (depicted in Fig 1), viral infection of target cells in the upper respiratory tract (URT) occurs with rate constant *β*. After spending an average time of 1/*k* in an eclipse phase *E*, infected cells enter a productively infected state *I*, where they produce virus at rate *p* (in the absence of N-R) and die at per capita rate *δ*. SARS-CoV-2 is cleared at per capita rate *c*.

**Fig 1.**
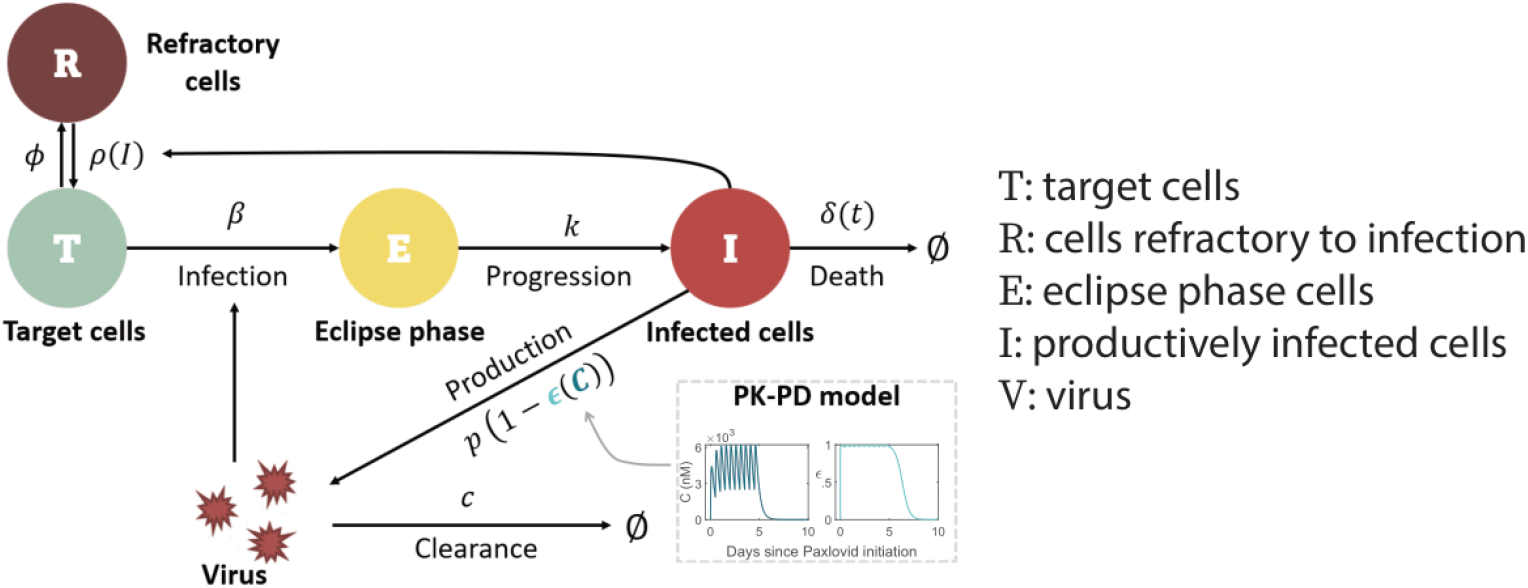
Schematic of the viral dynamic model. The model includes pharmacokinetic (PK) and pharmacodynamic (PD) sub-models, specifying how the drug concentration C and drug effectiveness *ϵ*(*C*) change over time (model details in Methods and S1 Text).

For the innate immune response, we assumed the amount of type-I and type-III interferons in the URT is proportional to the number of infected cells, *I*, and that interferon puts target cells into a temporary antiviral state (refractory to infection) [33,34,36–39] at rate *ϕ*.

Refractory cells become susceptible to infection again at rate 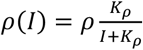, where *ρ* is the maximum rate at which refractory cells return to being susceptible [40] and *K*_*ρ*_ denotes the density of infected cells at which the rate of return is half-maximal^1^. Following Pawelek et al. [41], the adaptive immune response is modeled as causing an exponential increase of the death rate of infected cells (*δ*) at rate *σ* for a short time after its emergence time *t*^*^. This choice was motivated by the observation that virus-specific CD8^+^ T cells expand exponentially after viral infection [42]. This makes the death rate of infected cells a function of time *δ*(*t*). Finally, the concentration-dependent action of N-R is incorporated using a pharmacokinetic-pharmacodynamic (PK-PD) model. Additional details of the model formulation are provided in the Methods, S1 Text, and S1 Fig.

### Model describes the viral dynamics in all treated individuals

Our viral dynamic model describes the observed data for treated participants with and without rebound (Fig 2a). By fitting the model to the data, we obtain population (S1 Table in S2 Text) and individual (S2 Table in S2 Text) estimates of the model parameters, which are stratified by rebound vs. non-rebound (Fig 2b). The estimated time of infection relative to the time of symptom onset as reported by participants and the time of N-R initiation relative to infection and to symptom onset are also shown in Fig 2b. We found that the parameters (ρ, ϕ, *K*_ρ_) governing the dynamics of refractory cells, i.e., those cells that are protected from infection, are significantly different between individuals who rebound and those who do not. The differences in all of these parameters between the two groups were such that they favored the maintenance of cells in the refractory state in non-rebounders, who had a larger rate of cell entry into refractoriness *ϕ* (p=0.0004), a smaller maximum rate of cells returning to target status *ρ* (p=0.0047), and a smaller half-saturation constant for this process *K*_ρ_ (p=0.0056).

**Fig 2.**
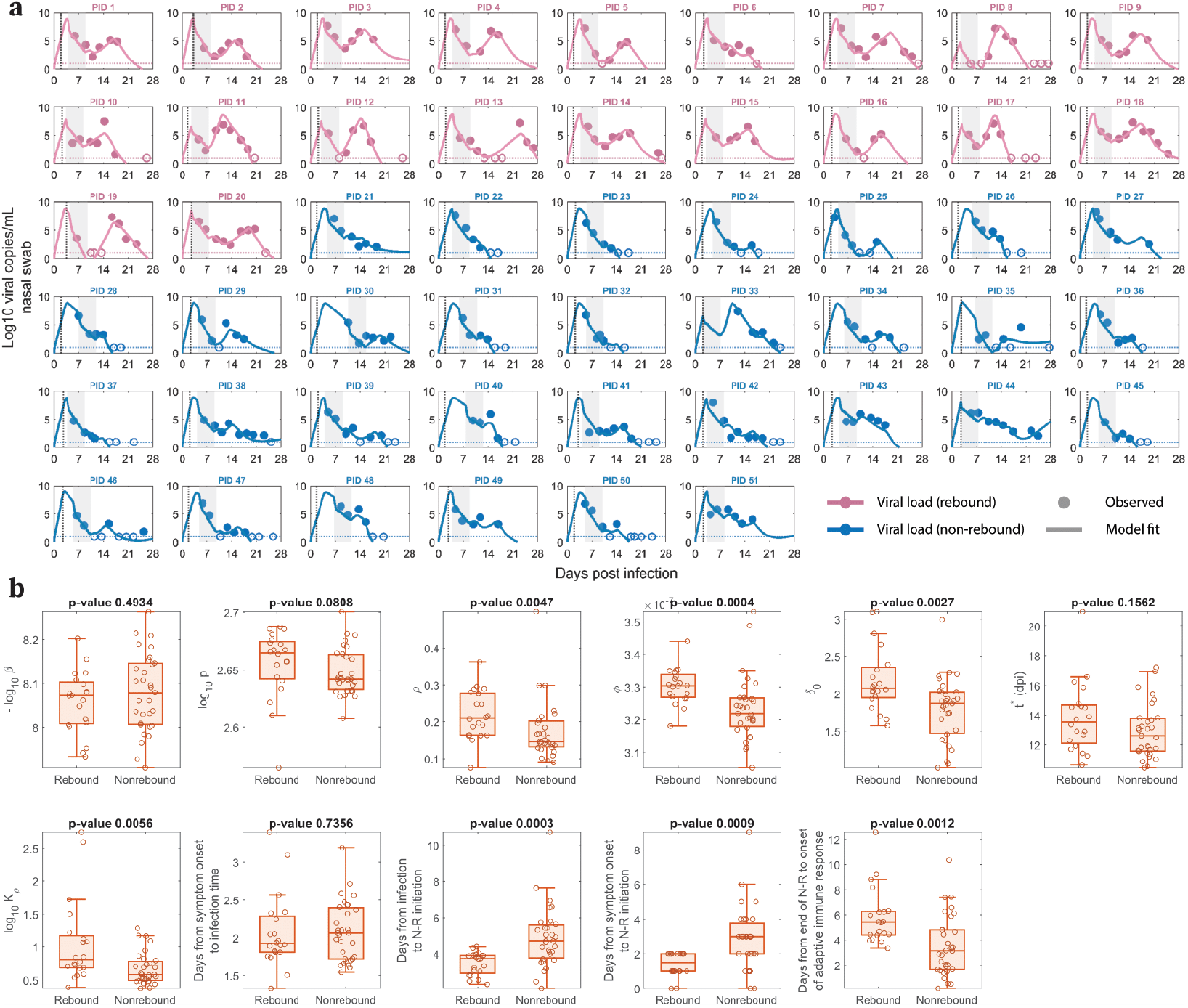
Model fits recapitulate viral dynamics and quantify differences in the characteristics between viral rebound and non-rebound individuals. **a**. Model fits to nasal viral loads of rebound (pink) and non-rebound (blue) individuals. The shaded area is the duration of N-R treatment. The dotted horizontal line is the limit of detection (LoD) for the RT-qPCR assay. Filled and open circles are data above and below the LoD, respectively. The dotted black vertical line indicates the reported time of symptom onset relative to the estimated time of infection. **b**. Box plots of best fit parameters and timing of N-R stratified by individuals who rebound vs. those who do not. The lower and upper limits of the box represent the first and third quartile, respectively. The line inside the box is the median and the whiskers connect the top/bottom of the box to the max/min values that are not outliers (data points further than 1.5 times the interquartile range). Overlaid circles are individual parameter values. Time of N-R initiation relative to symptom onset was recorded for each individual (except non-rebounder PID 23, whose symptom onset is imputed at one day prior to their first positive test). P-values are calculated using the Mann-Whitney U test.

In addition, the baseline infected cell death rate (*δ*_0_) was also significantly smaller in non-rebounders (p=0.0027). When we tested using “rebounder” as a covariate on each parameter to improve the model fit and to better understand factors distinguishing rebounders from non-rebounders, a covariate in *δ*_0_ provided the lowest BICc. However, the BICc difference was small (less than 4 points) compared to the model without a covariate (S3 Table in S3 Text).

Additionally, when we considered a variation of our best fit model with proliferation of target cells (details and model fit in S2a Fig in S4 Text), the baseline infected cell death rate was not significantly different between rebounders and non-rebounders (S2b Fig). On the other hand, there were still differences that are significant in the innate immune response parameters *ϕ* (p=0.0222) and *K*_ρ_ (p=0.0201). Specifically, in both models, the rebounders tend to have a larger value of *ϕ*, indicating a more rapid loss of target cells by going into the refractory state initially, and a larger value of *K*_ρ_, resulting in an earlier replenishment of target cells that can support viral rebound [43].

The time of N-R treatment relative to the estimated time of infection was about one day shorter in participants who rebounded vs. those who did not (median 3.75 days vs. 4.72 days, p=0.0003). This is consistent with the significant difference (p=0.0009) in the time of N-R initiation relative to the time of symptom onset in rebound vs. non-rebound individuals, as suggested before [6,27,43,44]. These differences in parameter estimates manifest in clear distinctions in model dynamics (viral load, target cells, infected cells) between rebounders and non-rebounders, as discussed and demonstrated in S3-4 Figs and S5 Text. A model variation that includes logistic proliferation of target cells discussed in S4 Text also predicts similar model dynamics (S5 Fig in S5 Text).

The model also recapitulates the data in untreated individuals from the same ongoing clinical cohort (S6a Fig in S6 Text). We also find that the parameter distribution between the treated and untreated groups are statistically similar (S6b Fig in S6 Text). The one exception is the average difference of 1.23 days (95% confidence interval [0.44, 2.03], p=0.0026) in the estimated onset time of the adaptive immune response, which is later in treated individuals compared to untreated individuals.

### The sensitivity of viral rebound to treatment initiation time and the duration of treatment

Our results suggest that the time of N-R treatment initiation and the availability of target cells at that time are critical to define whether a rebound occurs or not. To further explore this, we used simulation experiments to show that delaying or extending the period of treatment with N-R can decrease the probability of rebound. We simulated n=20 treatment cohorts, each with 100 randomly generated *in silico* individuals treated with N-R (see Methods for details), and assessed what percentage of individuals in each cohort exhibited rebound, defined as the viral load returning above 10^4^ RNA copies per mL [6]. Samples of the simulated viral dynamics for individuals in the *in silico* cohorts are presented in S7a-c Fig in S8 Text. Without treatment, our cohorts of *in silico* individuals have similar rebound statistics as those reported in the 8 clinical studies [6,17,19,20,23–26] (S7d Fig in S8 Text).

We tested treatment starting at days 1, 2, 3 and 4 post symptom onset, with symptom onset assumed to be 3 days post infection. Extending treatment could be a feasible method of preventing rebound [27,43,45], so we also examined a 5-, 6-, 7-, 8- and 10-day treatment courses. In one scenario, we assume N-R does not affect the development of adaptive immune response (Fig 3). In a second scenario, we assume that the onset of the adaptive immune response is delayed more with longer treatments (S8 Fig in S9 Text). It is important to examine this possibility as it would make rebound more likely. The time of symptom onset is fixed at 3 days post infection; however, assuming either 2 or 4 days does not change the general trend observed in Fig 3 and S8 Fig in S9 Text in which we observed a clear decrease in rebound percentage as treatment is initiated later. We also found that an increase in the duration of treatment with N-R tends to prevent viral rebound. In all scenarios, extending treatment to 10 days decreases the probability of rebound in our 20 simulated 100-person cohorts to a level so low that it does not occur for all practical purposes.

**Fig 3.**
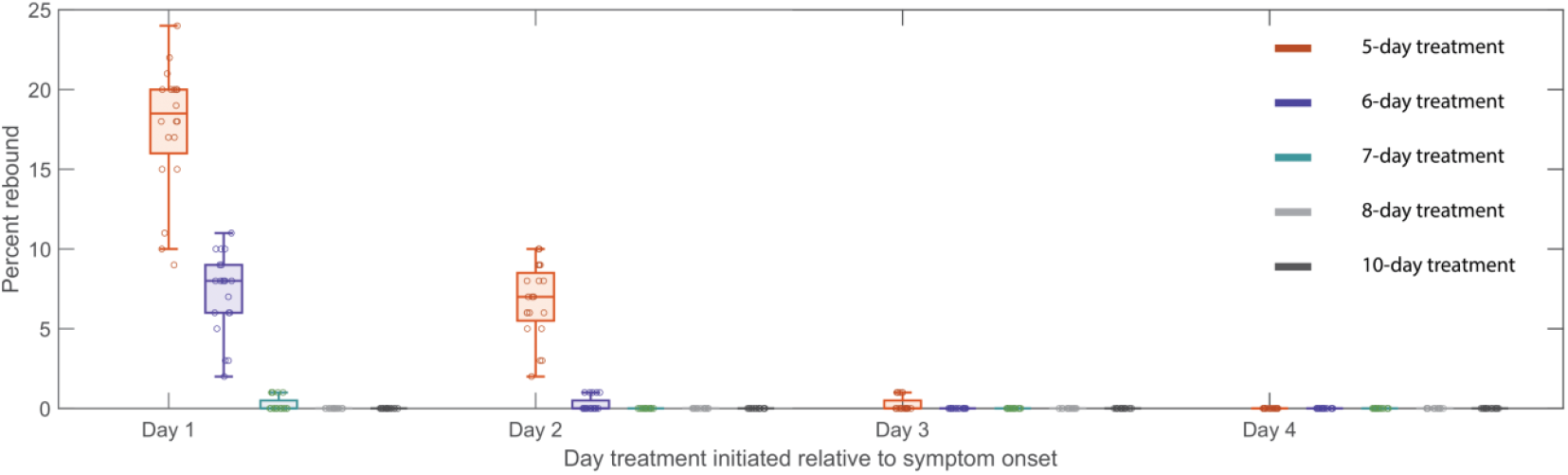
Predicted rebound relative to the time and duration of treatment. Predicted rebound relative to 5-, 6-, 7-, 8-, and 10-day course of N-R. Symptom onset is assumed to occur three days post infection. Boxplots depict the percentage of rebound cases from 20 *in silico* cohorts, each with 100 individuals, for different treatment initiation times. Each open circle represents the rebound percentage from one cohort. The extended duration of N-R (beyond a 5-day treatment course) is assumed to not cause additional delay on the onset of adaptive immune response.

## Discussion

Here, we extended a viral dynamic model of SARS-CoV-2 infection to show that the main driver of viral rebound in the setting of treatment is the preservation of target cells, often as a result of a robust innate immune response, or early treatment initiation. Our model shows that once N-R treatment is completed and the drug is washed out before an adaptive immune response develops, residual viable viruses can rebound if there are sufficient target cells remaining. Our results support our hypothesis [27] and echo the findings of recently published modeling studies [43,46]. However, our conclusions are supported by a more robust dataset of individuals treated with N-R, considerations of alternative models and assumptions on the impact of N-R on the development of an adaptive immune response with a detailed PK-PD model.

Our best model is able to capture the viral dynamics observed in all participants. It suggests that the protective effects of innate immunity preserved the majority of target cells by putting them into an antiviral state shortly after the virus starts growing exponentially (S3-4 Figs in S5 Text). During treatment, the viral load and the number of infected cells rapidly decline (Fig 2a and S4c, f Figs in S5 Text) due to infected cell death and continuous viral clearance, concurrent with reduced viral production due to drug activity. This decline leads to a decrease in the interferon response, causing cells to exit more quickly from the refractory state [36–40]. It is clear from the data of both rebound and non-rebound individuals that a five-day course of N-R is likely to be insufficient to completely eliminate the virus. Indeed, there was measurable virus (viral load > LoD) after the completion of treatment (the first data point after treatment) in 40 of the 51 participants (Fig 2a). Thus, if viable viruses remain after the drug is washed out and before an adaptive immune response can be mounted, virus can rebound. However, whether the virus rebounds to an observable level is also determined by the time between the end of treatment and the generation of an effective adaptive immune response, and to some degree, the differences in the maintenance of the cell refractory status (Fig 2b). This conclusion is supported by the observation that the time between the end of treatment and the predicted onset time of an adaptive immune response in the model is statistically different between the rebound and non-rebound groups. For the rebound group, the estimated time [min, max] is 5.87 [3.34, 12.56] days, and for the non-rebound group, it is 3.53 [0.14, 10.35] days (p = 0.0012) (Fig 2b). Note that this difference is not driven by the fitted onset time of the adaptive immune response *t*^*^ measured from the estimated time of infection, whose distribution is statistically similar between the two groups (Fig 2b). Instead, the difference in the time between the end of treatment and the onset time of the adaptive immune response is mainly driven by the earlier time of treatment initiation in the rebound group (Fig 2b).

The time of treatment initiation also plays a crucial role in determining if a rebound is observed or not. If treatment is initiated early after infection, before a time we denote *t*_*critical*_, a substantial number of target cells remain unprotected after the 5-day treatment and viral rebound is likely to occur. After *t*_*critical*_, too few target cells remain available to support viral growth; however, target cells still return from the refractory state as the virus is eliminated. Since viral growth switches to viral decay at the time of the viral peak in an untreated individual, this means *t*_*critical*_ is the time the viral peak is reached. In more technical terms *t*_*critical*_ corresponds to the time the effective reproductive number R equals 1, so that on average, each infected cell produces one new infected cell, leading to neither growth nor decay in the number of infected cells. In several observational/retrospective studies focusing on Omicron subvariants, the time to the viral peak is suggested to be 2 to 5 days post symptom onset [47–49]. We observed that for the participants in this study, who were all infected with Omicron subvariants, rebound is associated with treatment initiated within 2 days of symptom onset [6]. This suggests treatment might have been initiated prior to *t*_*critical*_ while the virus level is still expanding. Delaying treatment may be a strategy to reduce the possibility of viral rebound (Fig 3 and S8 Fig in S9 Text); however, delaying treatment could have a negative impact on the severity of disease in the high-risk individuals for whom N-R is recommended, and this question deserves more study [50]. In addition, N-R treatment accelerates viral clearance and hence potentially can reduce viral transmission. See Fig 4 for a summary description of our results.

**Fig 4.**
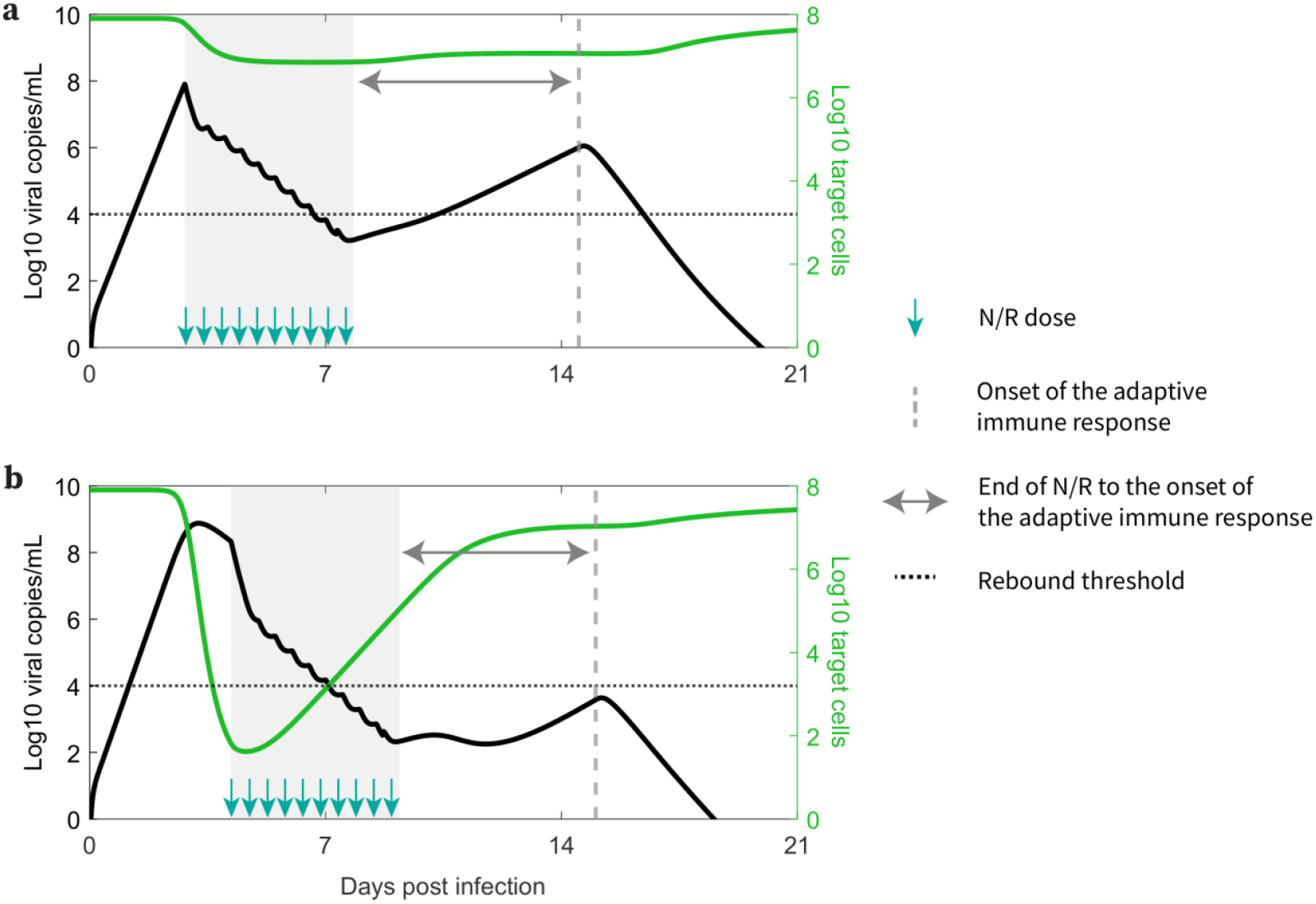
How early treatment correlates with higher rebound probability. **a**. Early treatments preserve more target cells and result in a longer duration between the end of N-R and the onset of an adaptive immune response, leading to a higher probability of an individual being classified as experiencing rebound. **b**. Later treatments preserve fewer target cells and result in a shorter duration between the end of N-R and the onset of an adaptive immune response, leading to a lower probability of an individual being classified as experiencing rebound.

Interestingly, all individuals studied here were vaccinated and boosted, and nonetheless had breakthrough infections with Omicron sub-variants [6]. Thus, while adaptive B and T cell immune responses did not prevent infection, they might have been present at the time of infection and could have affected the level of preserved target cells. The timing of the adaptive immune response and its expansion may play a crucial role in the occurrence of viral rebound. In particular, without a strong adaptive immune response, even a longer course of N-R still resulted in viral rebound in immunocompromised patients with severe disease [15,16,51]. Delaying the initiation of N-R may also provide more time for the priming of the adaptive immune response and shorten the time between the end of treatment and the emergence of the adaptive immune response, which would reduce the chance of rebound.

Our model predicted that the 20 rebound participants in the studied set have both innate and adaptive immune responses comparable to those of non-rebound participants (Fig 2b). This intriguing finding is supported by the clinical observations that most viral rebounds quickly resolve within several days [52] and this correlates with a strong antibody and T-cell immune response [13]. There is also contradictory evidence suggesting that N-R may delay the development of the adaptive immune response [53,54]. We found an average of 1.23 day delay in the estimated onset time of the adaptive immune response in the treated vs. untreated groups (S6 Text). Even so, the rebound participants quickly cleared the rebounding virus. This suggests that while early initiation of N-R may slightly delay the onset of the adaptive immune response, perhaps due to lower level of antigens, it does not stop the development of an adaptive immune response in non-immunocompromised individuals. Thus, if the adaptive immune response is not significantly impeded by treatment, prolonging treatment can be beneficial in reducing rebound and does not have the possible detrimental effects on disease severity or increase viral transmission of delaying treatment [44]. Indeed, using an *in silico* cohort we show that even a modest extension to a 6-day treatment course can significantly reduce viral rebound incidence (Fig 3 and S8 Fig in S9 Text). Extensions beyond a 6-day treatment course can further reduce rebound incidence with a 10-day treatment course almost totally eliminating the possibility of rebound in our *in silico* patient cohorts (Fig 3 and S8 Fig in S9 Text). A recent clinical trial compared 5 vs. 10 vs. 15 days treatment with N-R given to immunocompromised patients with COVID-19 (ClinicalTrials.gov: NCT05438602). The final analysis of 150 participants showed that extending treatment to 10 or 15 days can minimize the risk of rebound [55]. While 9 of 52 participants treated with 5 days of N-R rebounded, only 1 participant rebounded in the 10-day (n=48) and 15-day (n=50) treatment groups. While the clinical trial was carried out with immunocompromised patients, the single rebound incidence in the 10-day treated group supports our simulation results for a theoretical 10-day treatment for mild-to-moderate individuals with high risk of progression (Fig 3). When the cost of the drug is accounted for, the optimal treatment duration to minimize rebound and cost falls between 7 and 8 days (S9 Fig in S10 Text). However, because N-R is packaged as a 5-day course of treatment, extending treatment to 10 or 15 days may be more practical. Additionally, we previously suggested that the success of a second course of N-R once viral rebound occurs will also depend on the timing of an effective adaptive immune response in a similar manner [27]. This is corroborated by observations of recurring viral rebounds in an immunocompromised individual at the end of each treatment period, which eventually leads to the development of the resistance mutation E166V/L50F [15]. An ongoing clinical trial aims to investigate this possibility (ClinicalTrials.gov: NCT05567952).

Rather than extend treatment duration, the use of a drug with a longer half-life may be helpful, especially if infectious forms of SARS-CoV-2 can persist during antiviral treatment [8,9,56]. An ongoing clinical trial of ensitrelvir (ClinicalTrials.gov: NCT05305547) [57], a protease inhibitor that also targets SARS-CoV-2 3CLpro but with a longer half-life than nirmatrelvir [58], yielded results suggesting that this new drug was virologically active and did not significantly increase the risk of viral rebound [45].

The phenomenon of viral rebound has also been observed for monoclonal antibody treatments for SARS-CoV-2 [59–63]. One example is bamlanivimab, the first monoclonal antibody that received FDA emergency use authorization for the treatment of COVID-19 [61–63]. However, rebounds in the case of monoclonal antibodies are associated with the emergence of resistance mutations [59–63], which contrasts with the lack of evidence for resistant mutants *in vivo* in the majority of cases for the current antiviral treatments [5–7,11–13]. Yet, the emergence of resistance mutations to monoclonal antibodies does not always lead to viral rebound [59,60], suggesting other mechanisms beside selection pressure due to treatment may contribute to observable viral rebounds. Our previous modeling studies suggested that target cell regeneration mechanisms, such as homeostatic proliferation of epithelial cells [64–66] or refractory cells returning to a susceptible state, are necessary to explain the high amplitude viral rebounds observed in bamlanivimab treated participants [32]. Here our model with logistic proliferation (S4 Text) also recapitulates the viral load dynamics in rebound and non-rebound participants (S2a Fig in S4 Text) and the stratified parameter values also support the conclusion that early N-R initiations correlate with a higher probability of rebound (S2b Fig in S4 Text).

However, the net regeneration effect of target cells is similar to that in the innate immune response model (S5 Fig compared to S3 Fig in S5 Text). This is likely because potent target cell preservation limits the proliferation rate, which is related to the number of cells that are lost by infection. Moreover, because rebound occurs within days after the end of treatment, there is also not sufficient time for the proliferation effect to be more evident. In addition to explaining viral rebound, target cell regeneration mechanisms may also explain the observations of low amplitude viral rebounds/persistence in untreated individuals prior to the development of an effective adaptive immune response [67,68].

Our study has some limitations, the principal of which is not knowing the precise date of infection of each individual. This is a very common situation when dealing with infectious diseases [69,70], and it is ameliorated by using a well-established dynamical model, which in most cases allows us to infer the time of infection better than may be known clinically. Another important issue is that we do not have data on the immune response, even though we include both innate and acquired immune factors in our model. In the context of vaccinated individuals, this could be even more important, although it has been shown before that the viral dynamics of breakthrough infections maybe similar to that in unvaccinated individuals [71,72]. Our study could be strengthened and validated by incorporating detailed longitudinal immune response data, similar to those collected in the human challenge study for SARS-CoV-2 [73].

Furthermore, for the logistical proliferation model, markers of target cell proliferation or re-population could be used to support the model. We should also re-emphasize that although delaying treatment leads to lower probability of rebound, we do not evaluate the effect on severity of disease.

In summary, our results suggest the occurrence of viral rebound following a complete course of N-R may be due to the level of preserved target cells in the setting of incomplete elimination of the virus. Delaying initiation of treatment for a day or a few days following the first signs of infection should have some benefit in reducing the possibility of rebound, but at the cost of allowing viral growth to continue and the possibility of increased disease severity. On the other hand, extending treatments by several days may also reduce the likelihood of rebound, but at an increased cost of drug. We remark that viral rebound is not an intrinsic feature of our model, but rather a possibility within the model dynamical landscape. This is clearly demonstrated by the model fits to non-rebound individuals (treated and untreated). Lastly, rebound following antiviral treatments is not unique to N-R [21,28]. In particular, rebound without evidence of resistance has also been observed for the protease inhibitor simnotrelvir [28], which has a similar mechanism of action to nirmatrelvir and a shorter half-life [29]. Thus, these findings may provide an explanation for rebound following other antiviral treatments besides N-R.

## Methods Data

The data in this study comes from an ongoing observational cohort study. Full details of the study design and observations have been reported previously [6]. In summary, participants are adult outpatients selected from those who took part in the POSITIVES study (Post-vaccination Viral Characteristics Study) [7,74] within 5 days of an initial positive diagnostic test for COVID-19, had not yet completed a 5-day course of N-R, and had not received other antiviral or monoclonal antibody treatments [6]. Time of symptom onset was reported by participants and infection was confirmed with an initial PCR or rapid antigen test. Anterior nasal swabs were self-collected about three times a week for two weeks, then weekly until persistent undetectable results. The data were originally reported relative to the time of the initial diagnostic test [6]; however, we shifted the data to be “Days post infection” (Fig 3) based on fitting the model to the data (see Data Fitting). The primary definition for viral rebound was either (a) a positive viral culture following prior negative results, or (b) nadir viral load dropping below 4 log10 copies/mL then increased by at least 1 log10 copies/mL above the nadir and sustained above 4 log10 copies/mL for two consecutive measurements [6].

For this analysis, we selected all participants who took N-R and met two criteria: (1) had at least 5 data points, with (2) at least 4 of those data points above LOD. There were 51 participants that met these criteria (20 showing rebound and 31 showing no rebound).

Details regarding the statistics of rebound in untreated individuals are presented in S4 Table in S7 Text.

## Mathematical Model

We used an extension of the viral dynamic model, originally developed by Baccam et al. [75], Saenz et al. [76], and Pawelek et al. [41] to study acute influenza infections, which has previously been adapted to study SARS-CoV-2 infection dynamics [32–35]. The model below statistically outperformed the simpler versions used by Perelson et al. [27] (see S3 Table in S3 Text).

The model is described by the following set of ordinary differential equations:

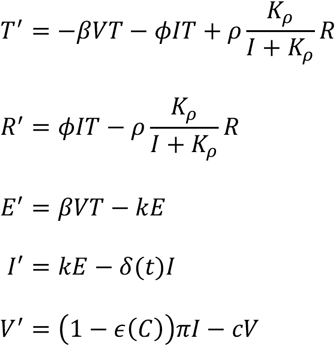

In this model, *T* is the number of target cells in the URT, *E* is the number of infected cells that have not yet started to produce virus, i.e., are in the eclipse phase, *I* is the number of productively infected cells, and *V* is the viral load. Target cells become infected with rate constant *β*. After being infected for an average time of 1/*k*, infected cells in the absence of therapy start producing virus at an adjusted rate *π* that accounts for sampling via a swab [33,34] and die at per capita rate *δ*, which we allow to be time dependent as described below. SARS-CoV-2 is cleared at per capita rate *c*.

For the innate immune response, we assume [34,41] the level of type-I and type-III interferons in the URT is proportional to the number of infected cells, *I*, because these cells produce IFN and recruit other IFN-producing cells, such as plasmacytoid dendritic cells. We also assume that interferon puts target cells in an antiviral state that is refractory to infection at rate *ϕ* [36–39]. The number of cells refractory to infection is denoted *R*. Refractory cells lose their protection and become susceptible to infection [40] at a rate 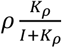. The density dependence of this rate on the number of infected cells *I* reflects the idea that when infected cells are abundant, they stimulate a strong interferon response, which keeps uninfected cells in a refractory state; but when infected cells decay below a critical threshold, they no longer sustain a sufficient interferon response to maintain cells in a refractory state and these cells return to being susceptible again [36–40]. Note that promoting a refractory state is just one possible mechanism of the innate immune system to fight SARS-CoV-2 infection [77]. A previous study by Ke et al. [34] examined various formulations (e.g., reduction in infection or viral production rate) that reflect different mechanisms of the innate immune response and found this formulation to be superior in capturing viral dynamics data.

We added to this model an adaptive immune response, since rebounds tend to occur late after infection, when adaptive immune responses have been observed [13]. As modeled by Pawelek et al. [41], we added this response to the model starting at time *t*^*^. We assumed that the adaptive response increases exponentially at rate *σ* for the short time period we model and causes an increase in the death rate of infected cells. This increased death rate could be due to the increasing presence of cytotoxic T cells or of viral-specific antibodies that bind to infected cells and cause their death by processes such as antibody-dependent cytotoxicity, antibody-dependent phagocytosis, or complement-mediated death. For simplicity, we fixed *σ* = 0.5 per day, which means that 1, 2, 3, 5 days after *t*^*^, the adaptive immune response will be at approximately 45%, 67%, 80%, and 93% of its maximum strength. The time-dependent infected cell death rate *δ*(*t*) takes the form:

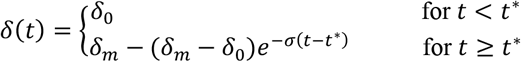

The effectiveness of nirmatrelvir in blocking viral replication and subsequent production of virions is given by 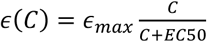, an E_max_ model [78] where *C* is the concentration of nirmatrelvir, *EC*_50_ is the concentration at which the drug effectiveness is half-maximal and *ϵ*_max_ is the maximum effectiveness. When *ϵ*(*C*) = 0 the drug has no effect and when *ϵ*(*C*) = 1 the drug is 100% effective at blocking virion production. Based on the complete model, viral growth occurs only when the fraction of remaining target cells is above a critical threshold, which is 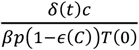, corresponding to the effective reproduction number R being larger than 1.

As it is impossible to know the number of viruses that initiated infection, we use a method suggested by Smith et al. [79] in which we assume the initiating virus is either cleared or rapidly infects cells. Thus, for initial conditions we use: *T*(0) = 8 × 10^7^ cells, *E*(0) = 1 cell, *I*(0) = 0, *V*(0) = 0, and *R*(0) = 0 as explained in Ke et al. [34]. They also noted that the infection dynamics are relatively insensitive to increasing the initial number of infected cells to 10.

### Pharmacokinetic and Pharmacodynamic Models for N-R

We assume the drug effectiveness *ϵ*(*C*) depends on the concentration of nirmatrelvir, *C*(*t*), according to an *E*_*max*_ model with EC50 = 62 nM, as presented in the FDA Emergency Use Authorization [1]. Following a *single dose* of 300 mg nirmatrelvir with 100 mg ritonavir, the observed maximum nirmatrelvir concentration is 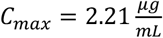. As nirmatrelvir has a molecular weight [80] of 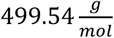 this value of *C*_*max*_ can also be expressed as 4.4 × 10^3^ *nM*. The half-life of nirmatrelvir when taken with ritonavir is about 6 hours [1], which corresponds to an elimination rate of 2.8/day. Additionally, dosing twice-daily achieved steady-state on day 2 with approximately 2-fold accumulation [1]. Using a simple multidose absorption-elimination model, the pharmacokinetics of nirmatrelvir is given by [78]

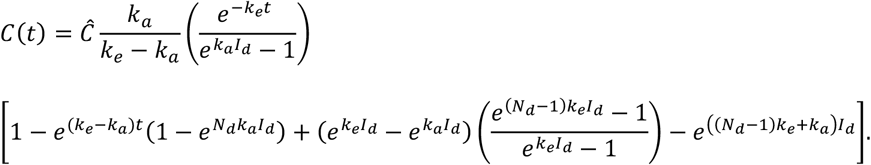

Here, *k*_*e*_ is the elimination rate (2.8/day), *k*_*a*_ is the absorption rate (17.5/day), *I*_*d*_ is the dosing interval (1/2 day), 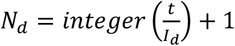 is the number of doses until time *t*, with the first dose at time *t* = 0. In S1 Text, we estimate 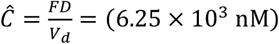. Details on the implementation of the pharmacokinetic model and the parameter values used can be found in S1 Text. With these assumptions, the drug effectiveness *ϵ*(*C*) hovers around 0.98 during treatment and then falls to zero rapidly after treatment stops (S1 Fig in S1 Text).

### Data Fitting

We used a nonlinear mixed effects modeling approach (software Monolix 2023R1, Lixoft, SA, Antony, France) to fit the model to viral load data for all 51 individuals simultaneously. We applied left censoring to data points under LOD.

We assumed that the parameters *p, δ*_0_, time of infection, and *K*_*ρ*_ follow a log-normal distribution. Parameters -log_10_ ϕ, -log_10_ *β, ρ*, and *t*^*^ were assumed to follow a logit-normal distribution, with ranges closely following literature values [33,34]. We constrained −log_10_ *β* between 7.5 and 9. Parameter *ρ* was constrained between 0 and 1 per day, − log_10_ *ϕ* between 5 and 12, and *t*^*^ between 7 and 28 days. No covariate was used during the initial fitting. A covariate based on whether a participant is classified as rebound or non-rebound was used later with the best fit model to determine the parameters that are different between these two groups.

The viral load data was originally reported relative to the number days since the initial PCR confirmation test. To estimate the time of infection, we shifted the data to be relative to the reported time of symptom onset. We then estimated the interval from the time of infection, or more precisely the time interval from when virus begins to grow exponentially as estimated by our model fitting, to when the participant reported symptoms. We then shifted the viral load data to be relative to this estimated time of infection.

The process to optimize the initial guesses of fitting parameters was done manually within the given parameter ranges to avoid unrealistic model dynamics. Whenever two models share a fitting parameter, the same initial guess for that parameter would be used in the fitting of both models. Model comparisons were done using the corrected Bayesian Information Criterion (BICc) [81] as reported by Monolix.

### Construction of an In-Silico Cohort

To quantify the chance of viral rebound after a five-day (or longer) course of treatment with N-R, we simulated a cohort of *in silico* patients. We used the following selection criteria to construct the cohort of *in silico* patients with typical viral load patterns: (1) The viral load must peak above 10^6^ copies per mL; (2) The peak must be reached between day 2 and day 7 after infection; (3) The viral load must decline below 10^2^ copies per mL by day 28. This algorithm is akin to a rejection algorithm, where we sample each parameter from the best fit population estimates (i.e., the estimated distribution) and only accept parameter sets that satisfy conditions (1) – (3). We fixed the time the adaptive immune response starts, *t*^*^, to the population estimate of 13 days, and set *δ*_*m*_ = 20/day to prevent unrealistic rebound once an effective immune response has been developed. Additional details of the *in silico* cohort are presented in S8 Text.

We used these admissible parameter sets to simulate treatment of different durations (5-, 6-, 7-, 8-, and 10-day of N-R) starting at different times (1 to 4 days post symptom onset) and calculate the probability of rebound. We also examined how a potential delay in the development of the adaptive immune response with longer treatment may affect the likelihood of rebound (S9 Text).

## Acknowledgements

The authors thank Jeremie Guedj for helpful comments on the manuscript.

## Funding

This work was performed under the auspices of the US Dept. of Energy under contract 89233218CNA000001 and supported by National Institutes of Health grant U54-HL143541-04 (RMR) Los Alamos National Laboratory LDRD 20200743ER (RMR), 20200695ER (ASP), 20210730ER (RMR), and 20220791PRD2 (TP).

Massachusetts Consortium on Pathogen Readiness (JZL, JEL, AKB, MJS) Massachusetts General Hospital Department of Medicine (JMV, AKB, MJS) National Institutes of Health grant U19 AI110818 (JZL, JEL, AKB, MJS) National Institutes of Health grant R01 AI176287 (JZL, JEL, AKB, MJS)

## Author contributions

Conceptualization: ASP, RMR, TP, JZL, MJS, AKB, JEL

Data curation: GEE, JB, RU, CM, MYL, MB, MCC, DT, KS, ZR, YL, SS, TDV, YK, JAS, SPH, ZW, JMV, JZL, MJS, AKB, JEL

Methodology: ASP, RMR, TP

Investigation: ASP, RMR, TP, JZL, MJS, AKB, JEL

Visualization: TP

Funding acquisition: ASP, RMR, TP, JMV, JZL, MJS, AKB, JEL

Project administration: ASP, RMR, JZL, MJS, AKB, JEL

Supervision: ASP, RMR, JZL, MJS, AKB, JEL

Writing – original draft: TP

Writing – review & editing: ASP, RMR, TP, GEE, JB, RU, CM, MYL, MB, MCC, DT, KS, ZR, YL, SS, TDV, YK, JAS, SPH, ZW, JMV, JZL, MJS, AKB, JEL

## Competing interests

ASP owns stock in Pfizer. He was also on a Pfizer advisory committee and received an honorarium. The other authors declare that they have no competing interests.

## Data and materials availability

The de-identified viral load data is provided in supplementary material (SM Data). Codes for fitting, plotting, and *in silico* analyses will be made available on GitHub once the manuscript is accepted and before its publication.

## Supplementary Materials

S1 to S10 Text

S1 to S9 Figs

S1 to S4 Tables

## Footnotes

1 Note if *I* ≫ *K*_*ρ*_, i.e. if the amount of interferon is very high, *ρ*(*I*) → 0, and cells remain in an antiviral state. However, as infection resolves and *I* becomes much less than *K*_*ρ*_, the antiviral state is lost at rate close to *ρ*.

